# Claws that snap: the raptorial mechanism of dryinid wasps

**DOI:** 10.64898/2025.12.19.695056

**Authors:** Brendon E. Boudinot, David Labonte, Adrian A. Smith, Sebastian Büsse, Jörg U. Hammel, Thomas van de Kamp, Carly M. Tribull

## Abstract

Prey capture is a key selection pressure, often favoring extreme performance morphologies. Many mechanisms of prey capture, however, remain poorly understood. In this study, we document the remarkable raptorial behavior and evolutionary biomechanics of the “pincer wasps” (Hymenoptera: Dryinidae). Dryinid claws, previously thought to function as simple vises (Nachtigall-&-Nachtigall-1974), are shown to possess “snap traps”, or claws that close rapidly upon prey contact. Detailed morphological analysis, using synchrotron-radiation microtomography (SR-µ-CT), reveals that all long-clawed dryinids can overcenter their specialized anterior pretarsal claws, presumably allowing elastic energy storage in the pretarsal apodemes. Uniquely, explosive release of this energy, leading to claw closure, appears to occur not through active withdrawal of a latch, but as the direct consequence of contact with the prey: impact hyperextends a specialized trigger-joint complex of the tarsus, pulling the claw back into undercenter alignment, and so freeing the stretched apodeme to fully recoil. Such a “contact trigger” is eminently useful, as these trap-claw wasps prey on insects with explosive jump capacity; by linking trap shutting with prey contact, success chances are increased. Paleontological and allometric analyses suggest that the raptorial behavior likely preceded the evolution of overcentering, which appears to have evolved via release from developmental constraint on anteroposterior symmetry, leading to a distinct scaling rule. These findings expand the documented diversity of spring-loaded biological mechanisms and establish Dryinidae as a promising system for studying how developmental constraints, mechanical demands, and functional innovation shape ecomorphological diversity.

**HIGHLIGHTS:**

- Dryinidae catch prey with spring-loaded trap claws, relying on a unique direct-action trigger.
- Raptorial behavior evolved first, followed by the functional derivation of two separate components.
- The evolution of novel raptorial mechanism was enabled by release of ancestral developmental constraint.

## BACKGROUND AND OBJECTIVES

The Dryinidae are a small but diverse family of stinging wasps with > 1,900 described species, which are globally distributed (Virla-et-al.-2023). These wasps are killer parasites (i.e., parasitoids) and predators that consume host and prey insects that use latch-mediated spring-actuated (LaMSA, Patek 2019, Longo-*et-al.-*2019) leg mechanisms to achieve extreme jumping performance: the leaf- and treehoppers (Auchenorrhyncha) (Burrows-2003, Burrows-et-al-2021, Olmi-1984, Virla-et-al.-2023). To grip the hosts or capture prey, they use the elongated claws of their prothoracic legs, which is a system that remains uncharacterized at the fine anatomical, mechanical, or behavioral-functional levels. Using high-speed videography (HSV), we first describe the behavioral and functional sequence of prey capture and assess whether the claws rely on direct muscle- or indirect spring-actuation. Next, using SR-µ-CT-based comparative anatomical observation of the leg system of Dryinidae and other Hymenoptera we identify the homologies of the system and the evolutionarily derived mechanical elements of the raptorial leg. Biomechanical analysis based on the HSV and SR-µ-CT comparisons then provides the foundation upon which we derive the biomechanical principles of the trap claw system. Finally, we assess the allometry of the functional anatomical elements of the system: the modified claw and its primary actuating muscle across a phylogenetically diverse sample.

## RESULTS AND DISCUSSION

### Functional morphology: The raptorial sequence

We captured the behavioral sequence of predation, identified conformational changes of dryinid external morphology, and estimated the rate of closure using high-speed videography (**Fig. 1**). The attack sequence begins when the dryinid has tracked and approached the victim (**Fig. 1A**), at which time both aroliar complexes (specialized adhesive organs) of the prothoracic pretarsi are planted on the substrate. The wasp then visually inspects the target and raises its prothoracic legs (**Fig. 1B**) before it begins to unfold the anterior pretarsal claws (**Fig. 1C**), popping into the fully opened position via a quick, jerky motion (**Fig. 1D**). The tarsi then slightly rotate toward the midline of the body (**Fig. 1E**) and, at the same time, the legs are slightly raised further, the tarsus bends at the joint between the third and fourth tarsomeres, and the wasp leans toward the target, elevating its abdomen in preparation for the initiation of the strike (**Fig. 1F**). The strike is initiated when the leg is whipped around the coxotrochanteral joint and lasts no more than ∼3.7 milliseconds from the beginning of the strike, with the legs folded, to the moment of chelar contact and claw closure (**Fig. 1G**). After successfully clamping on the body of the prey / host, the victim is then lifted, pulled towards the body, and stung (**Fig. 1H**), to be either parasitized through egg injection or consumed by the wasp. The clamping is so efficient that the wasp is able to maintain grip even against the extreme acceleration of the spring-loaded escape leap of the host (**Fig. 1K–M**). During the strike preparation, the joint between tarsomeres III and IV is bent and locked at an angle (**Fig. 1E’, F’**), but during the rapid strike itself, the tarsus bends at the joints of tarsomeres I and II (**Fig. 1I’, J’**).

**Figure 1.**
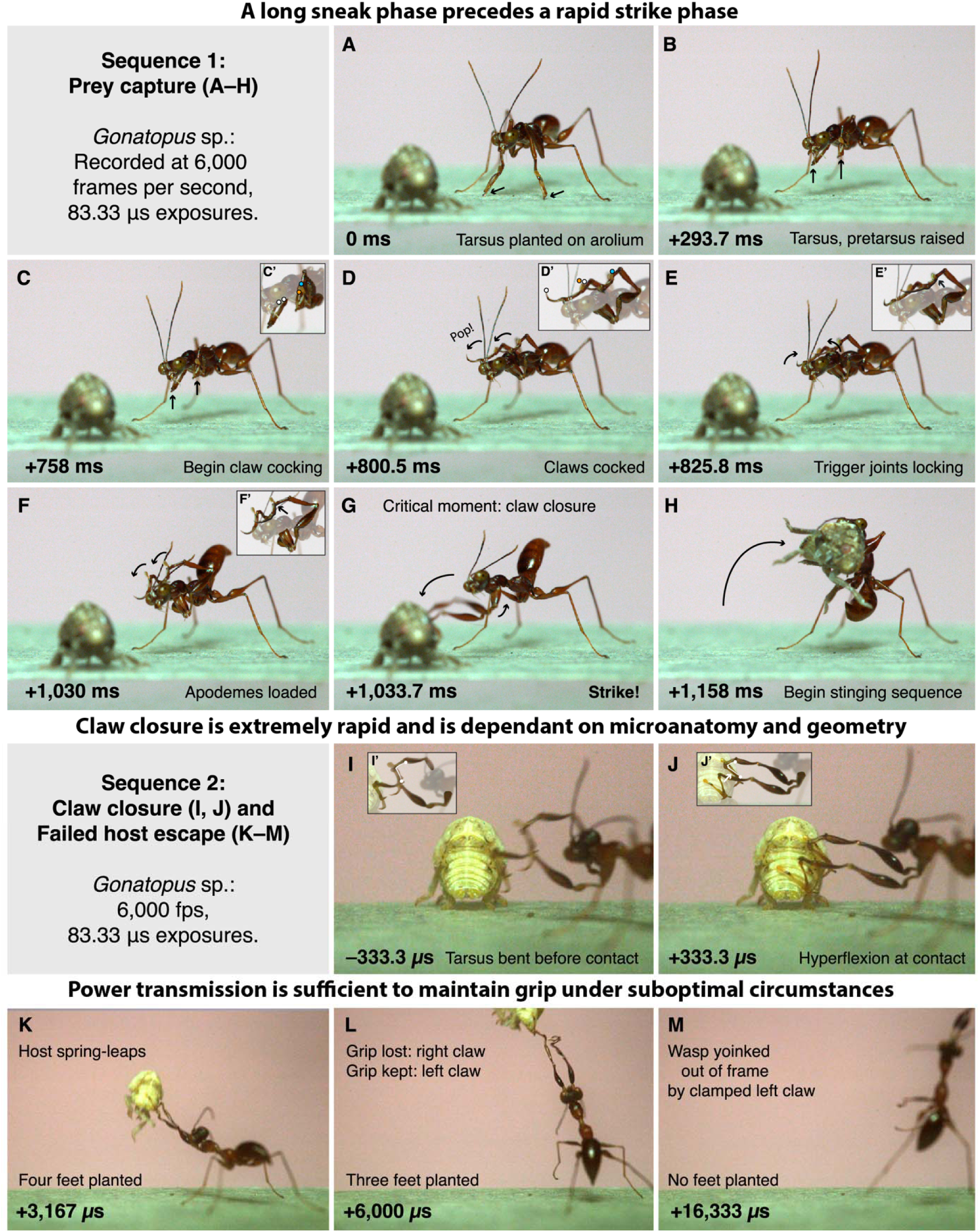
The behavioral sequence: High-speed video of *Gonatopus* strikes on planthopper nymphs. The first sequence shows a double-latch-bearing dryinid initiating and completing a trap claw attack. The second sequence shows claw closure and tarsal torsion (2a) and retention of grip through host escape leap (2b). Insets highlight claw cocking (**C’, D’**), and tarsal bending around the joints of tarsomeres I and II (**E’–F’**) and of III and IV (**I’–J’**). Both sequences presented in frames, with milliseconds (**A–H**) and microseconds (**I–M**) denoting elapsed time.

Claw closure for both tarsi took less than two exposures, each lasting 83.33 µs, thus took no longer than 166 µs. It appears that claw closure may result from physical contact with the prey itself, rather than by active neuromuscular actuation. Such a “contact-trigger” mechanism would appear sensible, for it will synchronize snapping of the trap with contact, improving efficiency of prey capture by avoiding premature or delayed closure. Using a congeneric species of the filmed dryinid as a proxy, we conservatively estimate that the power density (*i.e.*, mass-specific power) of the primary claw closer muscle was ≥ 808 W / kg (see Methods), in excess of the power density of striated muscle (*e.g.*, ∼ 400–500 W / kg; Josephson 1994, Pinot & Grappe 2014; Askew & Marsh 2002), This conservative estimate thus suggests that the claw may be spring-driven mechanism (Longo-*et*-*al.*-2019, Patek-2023), a hypothesis supported further by an anatomical analysis that reveals candidate structures that may act as latches (see below).

### The claw system: Groundplan and evolutionary morphology

To understand the physical causes of trap loading and snapping, we documented the anatomy and established the homologies and groundplan for the mechanisms used by a phylogenetically and phenotypically diverse sample of Dryinidae and outgroups. These include male and female aphelopine Dryinidae, which do not have the grasping mechanism (“achelate”), as well as the grasping-capable females (“chelate”) of the subfamilies Anteoninae, Thaumatodryininae, Gonatopodinae, and Dryininae. The raptorial system of female Dryinidae has been described as having a “chela” or grasping structure (XX_Olmi-*et-al.*-2020), comprising either: (a) an enlarged claw and a variably projecting fifth tarsomere, with an articulation between the two, as in females all living species (except Aphelopinae and Erwiniinae) and most fossil species; or (b) two claws, the arolium, and a lobe of the fifth tarsomere, as in the fossil, †*Raptodryinus*, from mid-Cretaceous Burmese amber (∼100 million years old), which has been assumed to be a mechanical intermediate.

From SR-µ-CT reconstructions of “chelate” Dryinidae (n = 10) and “achelate” Hymenoptera (n = 10), including achelate Dryinidae (two males, one aphelopine female), we observe that the enlarged claw of various female Dryinidae corresponds to the anterior pretarsal claw of male and achelate female Dryinidae and of other Hymenoptera (**Fig. 2A–C**), and that the articulation between the claw and fifth tarsomere is derived as a hinge in all sampled chelate individuals (**Fig. 2B, C**) as opposed to the monocondylic condition of achelate individuals (**Fig. 2A**). Further, †*Raptodryinus* represents a totally distinct mechanism, where grasping would have been affected through abduction of the arolium to a grossly enlarged plantar lobe of the fourth tarsomere (**Fig. 2D, E**)—rather than simply the “hinged lobe” as previously described XX_Olmi-*et-al.*-2020—implying that the raptorial behavior likely evolved prior to the origin of the trap claw mechanism of living Dryinidae. Loss of geometrical latching is implied for the ancestor of Aphelopinae, as the subfamily is nested within the phylogeny of the living Dryinidae (XX-Tribull-2015); female aphelopines appear to take advantage of hosts with side-stepping rather than leaping escape behavior (^7^-Virla-et-al.-2023).

**Figure 2.**
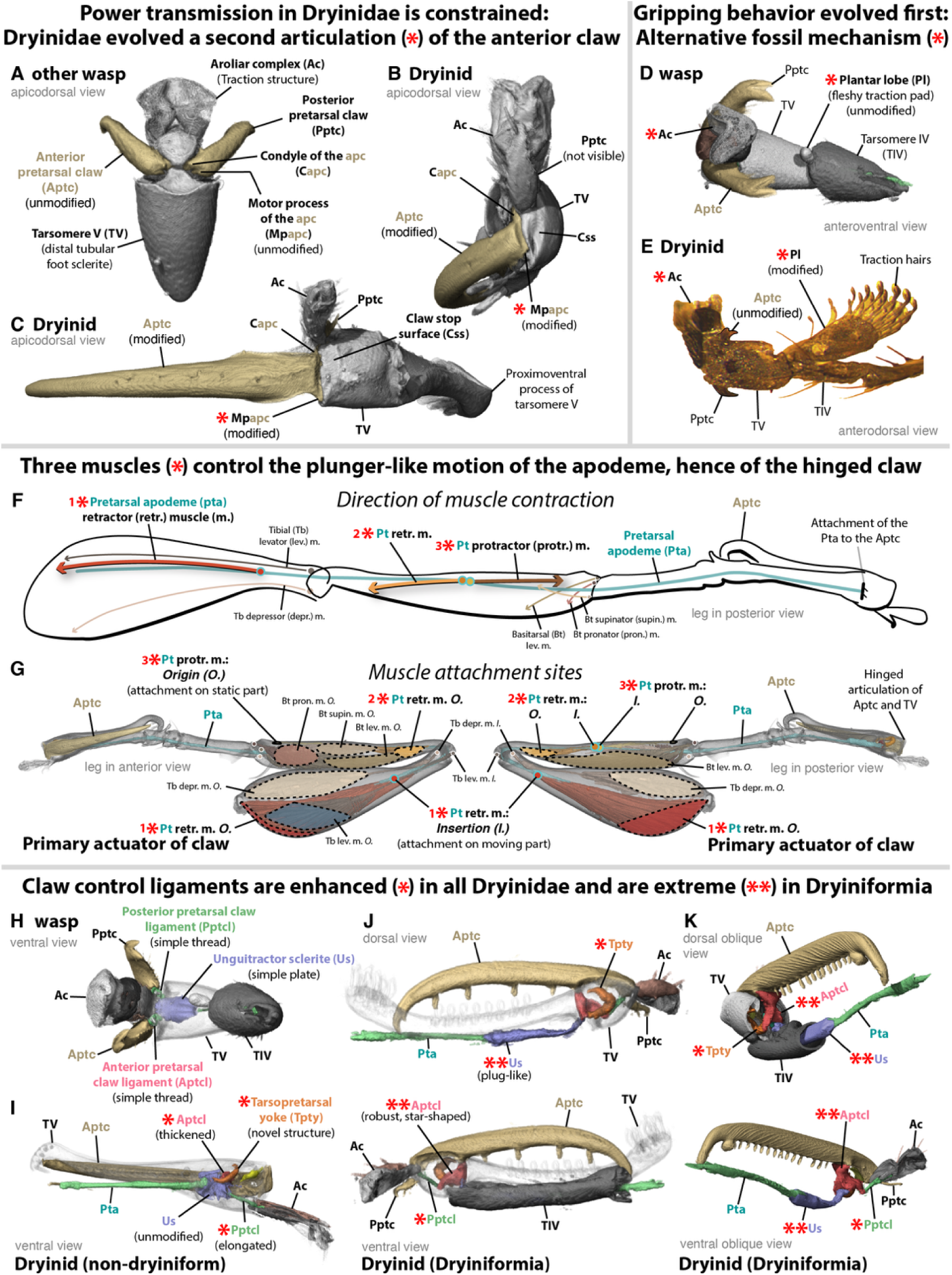
Comparative and functional morphology of the claw and leg system in Dryinidae. **(A–C)** Distal tarsomere and pretarsal complex in apicodorsal view. **(D, E)** Tarsal apex and pretarsal complex in anterodorsal and anteroventral oblique views. **(F)** Renders of prothoracic leg skeletomusculature distal to the trochanter; **(G)** pretarsal musculature schema from **(F)**. **(H–J’)** Unguitractor sclerite and distal ligament complexes in **(H, I, J’)** ventral view, **(J)** dorsal view, **(K)** proximodorsal oblique view, and **(K’)** proximoventral oblique view. *A, E, H = Trigonalys natalensis (BBKIT044); B = Anteon sp. (BBKIT071); C, F–G, L, M = Gonatopus mimoides (BBKIT093); D = †Raptodryinus, modified from Olmi et al. (2022); I = Anteoninae gen. et sp. indet. (BBKIT146); J = Echthrodelphax sp. (BBKIT055); K = Dryinus cf. sinicus (BBKIT028)*.

The claw system of Dryinidae and other Hymenoptera is a multisegmental complex that includes the femur, the tibia, the tarsus, and the pretarsus (**Fig. 2F, G**). The pretarsus consists of the arolium and its sclerites (see, *e.g.*, XX_Beutel-*et-al.*-2021, XX_Frantsevich-&-Gorb-2004, XX_Gladun-2008), the anteroposteriorly paired pretarsal claws, and the pretarsal apodeme and claw ligament complex. The apodeme is a tube-like invagination that extends from just ventrad the claws through the tarsus and tibia all the way into the femur; it bears a pair of ligaments that attach the motor processes of the claws to the unguitractor sclerite (**Fig. 2H**) (see also XX_Gorb-1995, XX_Heinzeller-et-al.-1989), as well as the genuflector sclerite in the femur, which is involved in contact latching in Orthoptera, flea beetles, and other taxa (XX_Burrows-1996, XX_Nadein-&-Betz-2016, XX_Földvári-et-al.-2019). In all sampled chelate Dryinidae, the anterior claw ligament is attached to the rim of the fifth tarsomere by a ligament-like structure that is coiled when the claw is closed (**Fig. 3E**) and uncoiled when the claw is open (**Fig. 3F**). This coiled ligament has no identifiable homologs among other Hymenoptera and appears involved in the overcentering mechanism as a safety factor preventing hyperextension; we hence name it the tarsopretarsal yoke (**Fig. 2I–K**). We further observe that the number of yokes varies from 1–3 across the Dryinidae, each of which can be homologized by their precise attachment locations. We refrain from defining these here.

**Figure 3.**
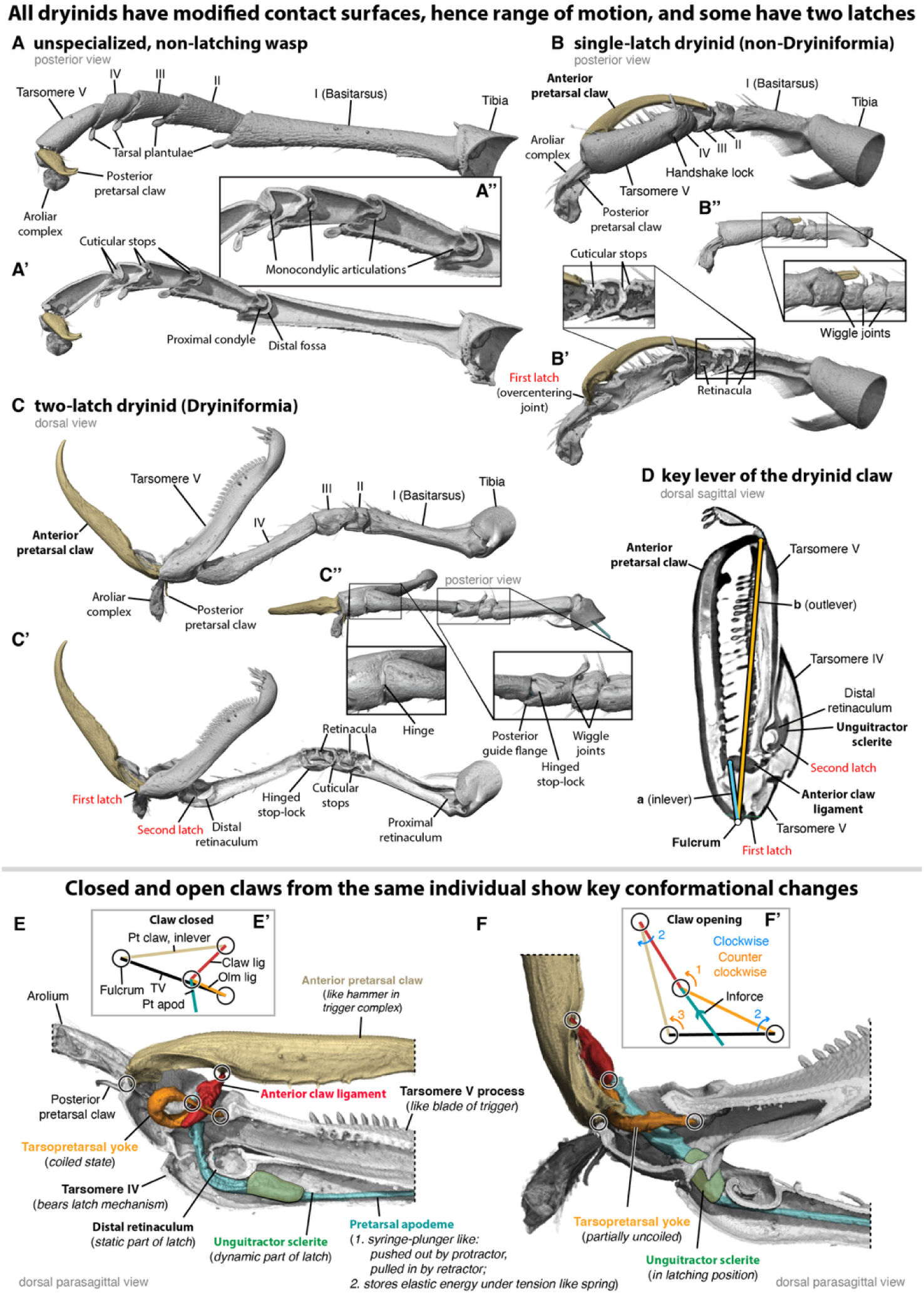
Tarsal anatomy, key lever, and four-bar mechanism of the raptorial claw. **(A, B)** Detail of the distal actuation complex of the clamping mechanism of an individual SR-µ-CT-scanned with one claw closed (A) and one partially opened (B). Insets **(A’, B’)** show four-bar mechanism abstracted from renders in (A) and (B). Impulse on the joint between the anterior claw ligament and pretarsal apodeme initiates torque (B’1), which propagates rotation around the joints between the anterior claw ligament and anterior pretarsal claw as well as of the tarsopretarsal yoke and tarsomere V (B’2), finally resulting in rotation about the proximal articulation of the anterior pretarsal claw and tarsomere v (B’3). Groundplan condition of Hymenoptera, as approximated by *Trigonalys natalensis* **(C–C’’)**. Geometric-latching-only form of raptorial mechanism and geometric-plus-contact latching mechanism, as shown by *Anteon* sp. (BBKIT071) **(D–D’’)** and *Gonatopus mimoides* (BBKIT093) **(E–E’’)**, respectively. **(F)** Planar mechanism of the anterior pretarsal claw and distal tarsomere mapped on a sagittal section of *G. mimoides*. *A, B, E, F = Gonatopus mimoides (BBKIT093); C = Trigonalys natalensis (BBKIT044); D = Anteon sp. (BBKIT071)*.

The pretarsal apodeme bears the muscles that actuate claw motion by moving the apodeme like a plunger proximad and distad, retracting and protracting the claws, respectively. The single protractor, *M. tibio-pretarsalis distalis* (Itipm2; newly recognized, *e.g.*, XX_von-Kéler-1955, XX_Aibekova-et-al.-2021_2025), originates distally on the tibia and is oriented proximally to its insertion on the apodeme (**Fig. 2F, G**). The tibial and femoral retractors of the pretarsal claws—*M. tibio-pretarsalis proximalis* (Itipm1) and *M. femoro-pretarsalis* (Ifpm1), respectively—have the opposite orientation, originating proximally and inserting distally (**Fig. 2F, G**). The femur has a dicondylic articulation with the tibia and bears the tibial levator, depressor, and chordotonal muscles (**Fig. 2F, G**). The femoral retractor appears grossly enlarged in “chelate” Dryinidae.

The tarsus is divided into five tarsomeres, but only the proximal tarsomere, the basitarsus, bears muscular insertions: those of the pronators, supinators, and levators of the basitarsus (**Fig. 2F, G**). The tibiotarsal joint and the four tarsotarsal articulations are ancestrally monocondylic, with a fossa-bearing process situated apicodorsally in the proximal sclerite and a condyle-bearing process proximodorsally in the distal sclerite (**Fig. 3A’, A’’**). The cone of motion of the tarsomeres is restrained by variably developed cuticular stops across Hymenoptera (**Fig. 3A’, A’’**). In Dryinidae, the proximal condyles of tarsomeres II–IV are reduced, the proximal rims are expanded, and the cuticular stops are modified, being shallow, roughly even, and circular rather than elliptical (**Fig. 3B’, B’’, C’, C’’**). In *Gonatopus*, the articulation between tarsomeres III and IV appears to form a hinged stop-lock with a guide flange (**Fig. 3C’’**). The joint between tarsomeres IV and V is differently modified, forming a rigid “handshake” lock as in *Anteon* (**Fig. 3B–B’’**) or a more mobile hinge as in *Gonatopus* (**Fig. 3C’’**). Across the majority of sampled Hymenoptera, tarsomeres I–IV were observed to have finger- or ring-like soft tissue that encircle the pretarsal apodeme. These retinacula, not previously recognized in the systematic morphological literature (*e.g.*, XX_Snodgrass-1935, XX_Beutel-et-al.-2014) or labeled but undefined (*e.g.*, “fixators” of Frantsevich-&-Gorb-2004), are broadly analogous to the ligamentous inferior and superior extensor retinacula of the ankle (Kapandji-2011) (*i.e.*, human tibiotarsal joint); they likely control the moment arms around the joints during muscle contraction by preventing bowstringing. We observed a proximal and distal retinaculum in the basitarsus of most taxa, a single and variably situated retinaculum in tarsomeres II and III, and a distal retinaculum in tarsomere IV (**Fig. 3B’, C’**).

### Biomechanical hypotheses for the trap claw system

We propose a set of mechanical hypotheses for the actuation of the trap claw (Figs. 3, 4), extracted from the combination of the behavioral and conformational sequences recorded in the high-speed videography (**Fig. 1**), our anatomical observations (**Figs. 2, 3**), and mechanical calculations (see Methods). Specifically posit that: (1) The basic mechanism for loading strain energy into the spring-like pretarsal apodeme among chelate Dryinidae is overcentering of the pretarsal claw inlever relative to the anchor point of the tarsopretarsal yoke (**Fig. 3E, E’, F, F’)**, forming a geometrical latch (**Fig. 4C, C’, D, D’**) (XX_Longo-et-al.-2019, XX_Steinhardt-et-al._2021), buttressed by robust cuticular stops (**Fig. 2B, C**) and further prevented from hyperextension by the yoke; (2) the basic latch release mechanism is a unique direct-action trigger formed by the modified joints of tarsomeres II and III (**Fig. 4F**), around which the fully taut pretarsal apodeme is forcibly jerked when the strike hyperflexes these articulations (**Fig. 1I, J**); and (3) in a subset of Dryinidae (Dryiniformia nom. nov. for clade Dryininae + Gonatopodinae + Thaumatodryininae, XX-Tribull-2015, XX-Martins-&-Melo-2024), the power output is enhanced by the evolutionary derivation of a contact latch mechanism between the pretarsal apodeme and the distal retinaculum, which is modified as a bulge-like invagination (**Fig. 3E, F**), reminiscent of “Heitler’s lump” in Orthoptera and homologous bulges in other taxa (XX_Burrows-1996, XX_Földvári-et-al.-2019).

**Figure 4.**
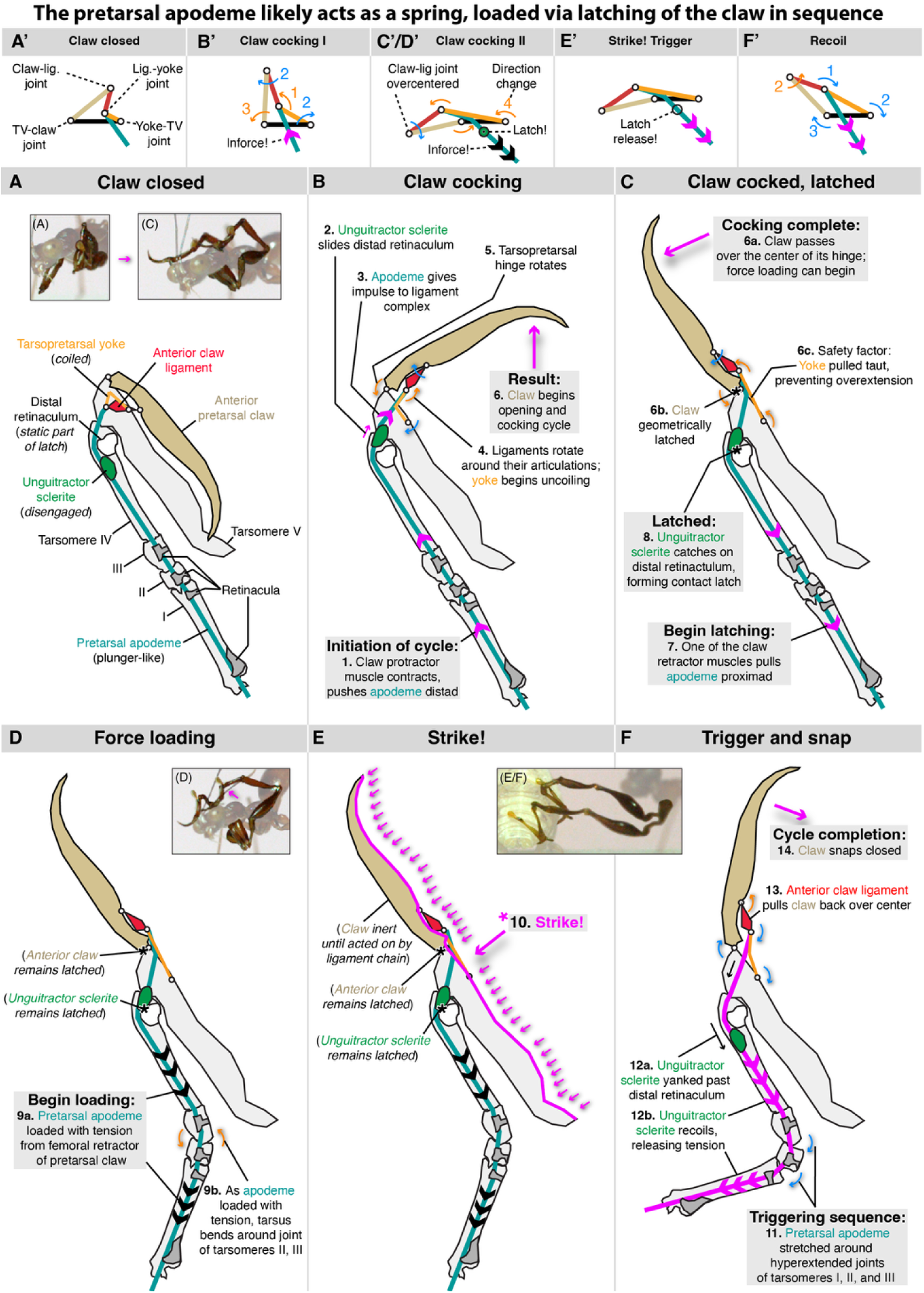
Key frames of the claw cycle and the hypothesized latching mechanism. **(A–F)** Diagrammatic presentation of mechanical hypothesis for the double-latched morphology observed in *Gonatopus* and other Dryiniformia, based on anatomical architecture, abstraction of the four-bar mechanism from the specimen scanned with one claw open and one closed, and select key frames from the high-speed videography. **(A’–F’)** Four-bar mechanism sequence. *Note: Morphological diagrams artificially separate the distal tarsomere from tarsomere IV to ease labeling; see the behavioral sequence for conformation in vivo. Model species: Gonatopus mimoides (BBKIT093)*.

In female Dryiniformia, catching of the unguitractor sclerite on the distal retinaculum is enabled by elongation of the pretarsal apodeme distal to the sclerite (**Fig. 3E, F**). This results not only in the ability of the unguitractor sclerite to be drawn into the housing of the fourth tarsomere (**Fig. 2J, J’** vs. **2H, I**), but it is also associated with complete rearrangement of the distal ligament complex. In Dryiniformia, the unguitractor sclerite is no longer the hub of ligamental attachment. Rather, the anterior pretarsal claw ligament is grossly modified, having a robust hour-glass shape and bearing the attachments of the tarsopretarsal yoke and the posterior claw ligament, in addition to the pretarsal apodeme (**Fig. 2I–K, 3E, F**). This modification may be adaptive, potentially preventing destruction of the ligament complex under increased strain due to gain of the tarsopretarsal contact latch, as compared to the unguitractor-as-hub conformation of the anteonine dryinids and other Hymenoptera. Whether or not the anterior-claw-ligament-as-hub organization is an adaptive response, its occurrence is directly associated with that of the contact latch configuration of the unguitractor sclerite and distal retinaculum, providing circumstantial evidence that the gain of contact latching enhanced the achievable power output succeeding the origin of geometric latching.

Several structural and functional elements of the raptorial fore leg of female Dryinidae are unique in comparison to other Arthropoda, beyond the ligamental modifications in the double-latched Dryiniformia. Among raptorial insects, the tarsopretarsal snapping and clamping mechanism of Dryinidae is distinct, as other groups—such as mantises, mantispid lacewings, *Ochthera* flies, and various predatory Hemiptera—use the femorotibial joint and its musculature for grasping prey (XX-Bäumler-*et-al.-*2023_XX-Gorb-1995_XX-Földvari-*et-al*.*-*2019). A possible exception is Enicocephalidae (Hemiptera), but descriptions of hunting behavior are limited and their tarsopretarsal morphology suggests static gripping (XX-Molleman-&-Walter-2001), rather than the dynamic snapping observed here. Similarly, lice have a tarsopretarsal gripping modifications, but these are used for locomotion rather than prey capture (XX-Preuss *et al*. 2024). Among arthropods with spring-loaded mechanisms, Dryinidae appear unique in that their trigger mechanism appears to act directly on the overcentered claw or tarsopretarsal catch via impact forces, delivered as a result of the strike. In contrast, the physical triggering of latch release in other groups either relies on muscular action (mantis shrimps XX_Patek-2019_2023, grasshoppers XX_Burrows-1996, trap jaw ants XX_Gronenberg-1996, XX_Booher-et-al.-2021), or depends on the deformation and slip of the striking structure itself for acceleration and force transfer (snap-jaw ants XX_Gronenberg-et-al.-1998, XX_Larabee-et-al.-2018 and termites XX_Seid-et-al.-2008). The exact anatomical trigger mechanism of Auchenorrhyncha, the hosts of Dryinidae, remains unaccounted for (XX_Burrows_2006_2007, XX_Burrows-et-al.-2021, XX_Siwanowicz-&-Burrows-2017). From a behavioral and ecological perspective, however, the host triggering mechanism is not specifically relevant, as dryinids may take advantage of the latency between the “PADT” (pleural-arch–depressor-trochanter) loading mechanism of Auchenorrhyncha (XX_Burrows-et-al.-2021) by their sneaking behavior (**Fig. 1**) and the use of geometric latching across all chelate species (**Figs. 2B, C, 4**), coupled with contact latching in Dryiniformia (**Fig. 3E, F**).

### Evolutionary allometry

We recognize the developmental rule changes for claw growth, mechanical advantage, and musculature that define the Dryinidae as a monophyletic group with specialized raptorial abilities. To do so, we used our SR-µ-CT data to extract femorotibial knee width (**W_tib_**) as a proxy for body size, the moment arms of the clamping mechanism (**Fig. 5A–C**), the mechanical advantage (MA, **Fig. 5D**), and the physiological cross-sectional area (PCSA) of the femoral claw retractor based on the preserved geometry of the muscle, as geared by the MA of the claw (**A_eff_** = **MA** • **PSCA** (**V_m_** [muscle volume] **• cos(P°)** [pennation angle] **/ L_f_**. [muscle fiber length]); XX-Püffel-et-al-2021). We found that, while the allometric slopes were not statistically different between chelate and achelate individuals, the intercepts of all chelate dryinid measurements differed significantly (**Fig. 5**). The MA of the claw in chelate Dryinidae is significantly lower than that of achelate dryinids and other Hymenoptera (**Fig. 5C**), but **A_eff_** was greater in chelates (**Fig. 5D**), indicating greater capacity for force transmission.

**Figure 5.**
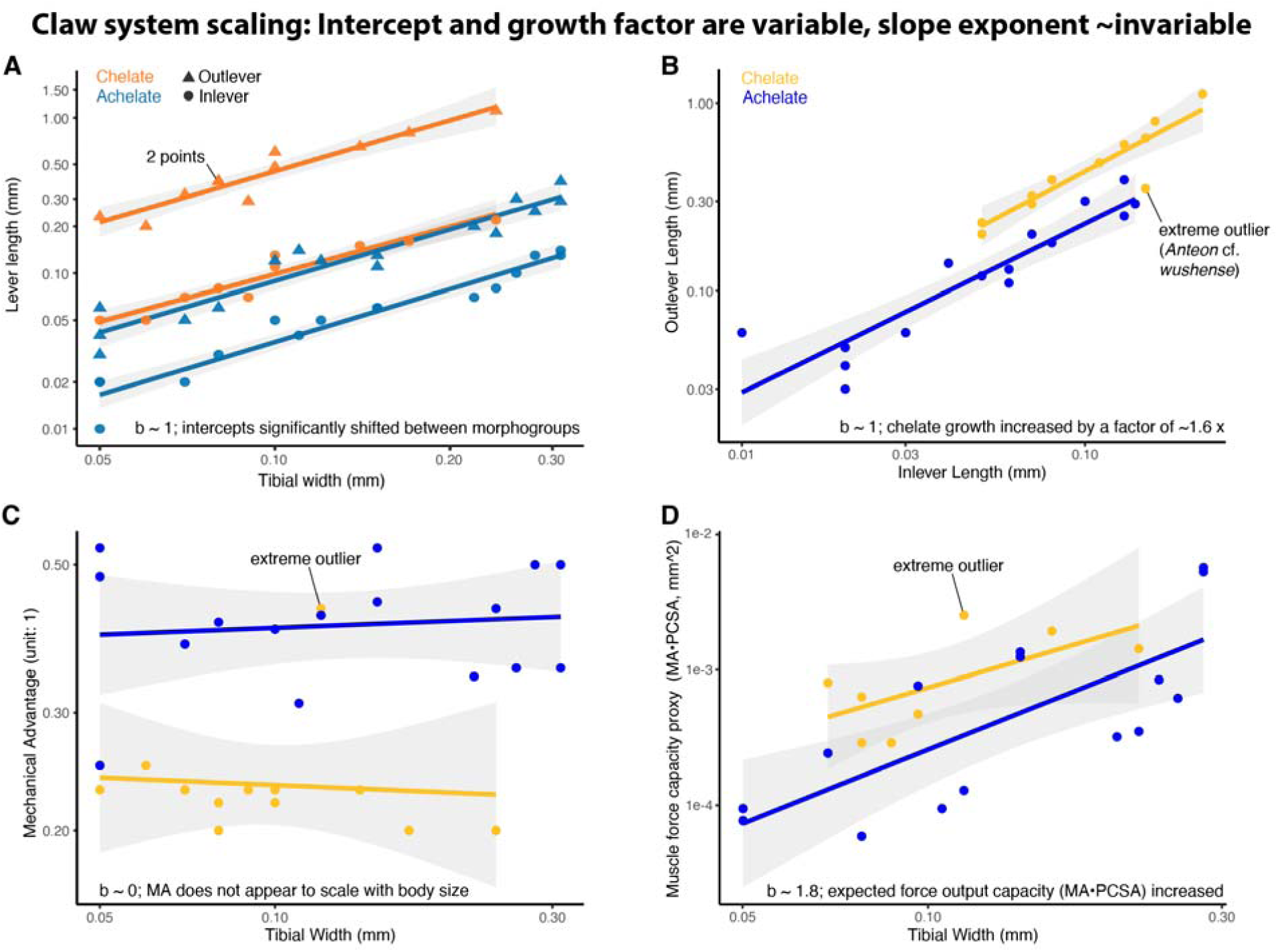
Evolutionary allometry of the anterior pretarsal claw and its actuating muscle. Allometric relationships of key functional elements of the claw mechanism in chelate Dryinidae versus achelate Dryinidae and other Hymenoptera, calculated using ordinary least squares (OLS) and with log scaled data plotted on raw axes. **(A)** Inlever (*a*) and outlever (*b*) length against tibial width, the characteristic length for the leg system. The slopes for both levers do not differ between the long- and short-clawed morphogroups (p_inlever_ = 0.438, p_outlever_ = 0.959); they are not significantly different from isometry (p_inlever_ = 0.085, p_outlever_ = 0.241) and are well explained by the model (R^2^_inlever_ = 0.945, R^2^_outlever_ = 0.954). The intercept of the long-clawed group is significantly higher (inlever: + 0.321, p = 0.040; outlever: + 0.696, p < 0.001). **(B)** Mechanical advantage plotted as outlever length (*b*) against inlever length (*a*). The slopes (R^2^ = 0.933) do not differ between the long- and short-clawed morphogroups (p > 0.287) and although somewhat hypoallometric (m = 0.90) are not significantly different from isometry (p = 0.40). Heteroscedasticity-consistent (HC) tests indicate that the intercept of the long-clawed group is significantly (p < 0.05) to marginally higher (p < 0.10) than that of the short-clawed group. Independent of body size, chelate Dryinidae have outlevers that are ∼1.6 x as long as achelate Dryinidae and other Hymenoptera. **(C)** Mechanical advantage does not scale with tibial width in either morphogroup (achelate slope = 0.034, p = 0.61; chelate slope ≈ −0.10, p = 0.41), and group-specific slopes did not significantly differ (interaction p = 0.33–0.75 across HC1–HC4 robust estimators). Isometry was strongly rejected (p < 10[¹²), indicating size invariance of MA. The chelate morphogroup showed a pronounced negative intercept shift (β ≈ –0.26 to –0.39, p < 0.001 under HC robust SEs), reflecting consistently lower MA at all body sizes. MA is thus a size-independent trait in all sampled Hymenoptera, with morphological differentiation between chelate and achelate morphogroups arising from an intercept shift rather than divergent scaling regimes. **(D)** The expected force output capacity (Aeff = physiological cross-sectional area (PSCA: V_m_ • cos(θ) • Lf^-1^) • mechanical advantage (MA)) of the major pretarsal claw closer muscle (Ifpm1) shows strong positive allometry with tibial width (log-log slope ∼ 1.8, p << 0.001), significantly exceeding isometry (test of slope = 1, p ∼ 0.046). The slopes were not significantly different between long- and short-clawed morphogroups (p > 0.5) but the chelate group had a positive intercept shift (Δlog Aeff ≈ 0.48, p < 0.01 under HC-robust SEs), corresponding to ∼1.6-fold higher Aeff at a given body size.

The finding that the intercept of the allometric slope is labile but not the slope itself is consistent with experimental evolutionary studies on allometry (XX-Pélabon-*et-al.-*2014, XX-Voje-*et-al.-*2014). This implies that the intercept shift is an adaptive response to directional natural selection on claw length during the early evolution of the Dryinidae (> 100 million years ago, XX-Martins-&-Melo-2024), with subsequent stabilizing selection, excepting an outlier group with exceptionally long inlevers (represented here by *Anteon* cf. *wushense*, **Fig. 6**). As the anterior claw is disproportionally long relative to the posterior claw, and as both claws are similar or roughly similar in size across Hymenoptera, the breaking of developmental constraint on the claw symmetry, is likely the consequence of changes in patterning of the anterior and posterior compartments of the prothoracic leg primordia during embryogenesis. Such changes also involved the gain of a second proximal articulation of the claw, present in all chelate species, followed by the derivation of a likely friction latch in the ancestor of the Dryiniformia nov. nov., potentially enabling even greater load on the spring-like pretarsal apodeme. The non-dryiniform lineages (e.g., Anteoninae) need to be videographically documented. In either case, the allometries of the anterior claw do not constrain the evolution of its functional surfaces, which display extremely taxonomically useful variation, i.e., diagnostic conditions of the claw armature such as teeth, and of the presence of broadened, spatulate hairs, among other character sets. Moreover, while the system appears to be under strong stabilizing selection, an extreme outlier species, *Anteon* cf. *wushense*, exhibits a claw with a proportionally long inlever, indicating that the gearing of the claw is still a labile trait, hence inlever length to body size is not under constraint of their allometric relationship.

**Figure 6.**
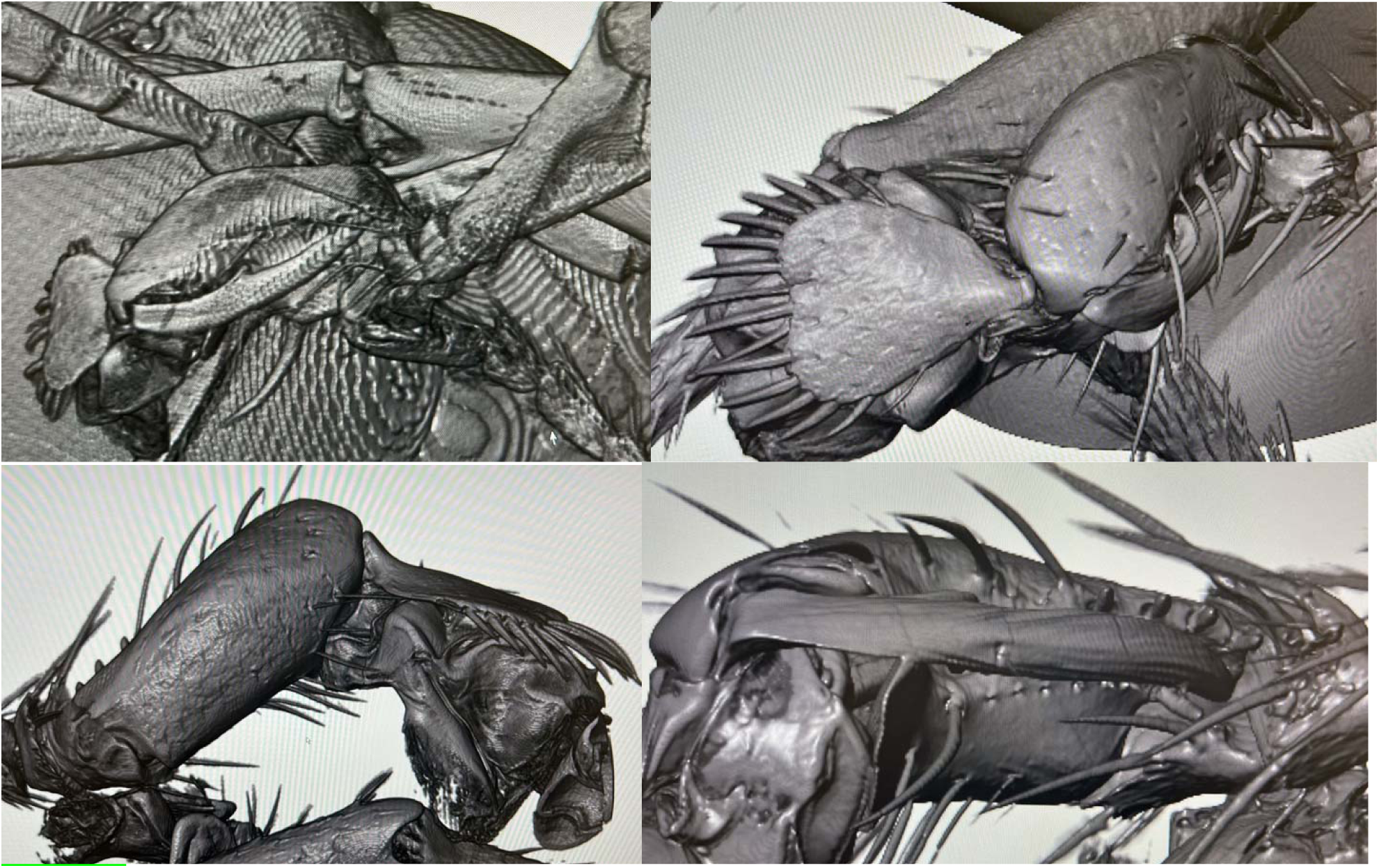
An outlier dryinid indicates existence of a second adaptive peak for chelates. The species *Anteon* cf. *wushense* is an extreme outlier among the long-clawed (chelate) Dryinidae. This species has a proportionally very long inlever while maintaining a comparatively strong claw actuator muscle, indicating that it has exceptionally strong grasp among all sampled Hymenoptera. Comparison of the anatomy of *Anteon* cf. *wushense* against other *Anteon* species and against Dryiniformia shows that this species also has an extremely large aroliar complex. The cause of the change in inlever length remains unclear.

The functional optimization of the raptorial mechanism appears to have been achieved through an increase in growth factor of the anterior claw, resulting in slope shifts for the lever arms and mechanical advantage. From a ‘design’ perspective, it is notable that the optimization has been achieved without sacrificing walking ability (**Fig. S4**) and that, relative to the chelae of crabs, for example, the primary actuating muscle is not within the housing of the claw and its articulating part. This may enable the modular evolution of the system, allow allowing the major closer muscle of the claw to become massive, while the claw and the tarsus itself remain light. Such a dynamic is not possible for crabs, where the closer muscle is within the mechanical housing of the chela, thus imposing a physical limitation on the co-optimization of power and rapid mobility of the claw as increase in the PCSA of the closer muscle necessarily increases the mass of the claw itself. In contrast, with a light and powerful claw, the wasps are able to whip their snap-traps onto their prey and hosts. Another analog may be the spearer-morph mantis shrimp, but their victims are impaled, rendering them useless for raising young. The trap claw system of Dryinidae, therefore, represents a unique solution to both prey capture and brood care.

## CONCLUSION

Our findings establish Dryinidae as an evolutionary biomechanical study system with a unique life history, morphological configuration, and trigger mechanism. High-speed videography coupled with synchrotron microtomography (SR-µ-CT) data suggest that at least some Dryinidae have a spring-loaded claw closure mechanism, leading us to conclude that these animals are snap claw wasps. We used SR-µ-CT to conduct a systematic survey of skeletomuscular anatomy across outgroups and the chelate and achelate phenotypes of Dryinidae, allowing us to resolve the homologies of the anatomical system. Based on our videography and SR-µ-CT data, particularly two individuals with one open and one closed claw, we propose explicit mechanical hypotheses for the opening and closing cycle and identify potential geometric and contact latches in the tarsopretarsal complex. Through comparison of fossils and a phylogenetically diverse sample of Dryinidae, we were further able to trace the evolution of key components across phylogeny, which indicates a sequence of behavioral derivation, followed by the gain of geometrical and contact latching in sequence. Our allometric analysis further shows that the clamp mechanism is highly constrained with one notable exception, whereas there is a potential trade-off escape in small individuals, linked to the out-of-housing configuration of the actuating muscles. Overall, our work enables the use of Dryinidae to systematically probe the origins of developmental and functional morphological novelty and the dynamics of force-transmitting complexes through deep time.

## ACKNOWLEDGMENTS

We are grateful to Martin Hauser, Minsoo Dong, and Daniel Tröger for sharing specimens for SR-µ-CT scanning, and Lisa Tewksbury, Dana Terrill, Dan Farnworth, and Alexandra Johnson from the University of Rhode Island for rearing and sharing the wasp used for high-speed filming. For comments on the pre-submission draft, we are grateful to Rolf Beutel and Ziv Lieberman. For encouragement and insightful discussion about dryinid natural history, we thank Adalgisa Guglielmino. We acknowledge the KIT Light Source for provision of instruments at their beamlines, and we would like to thank the Institute for Beam Physics and Technology (IBPT) for the operation of the storage ring, the Karlsruhe Research Accelerator (KARA). We thank Tomáš Faragó for tomographic reconstruction and Angelica Cecilia and Marcus Zuber for their support during beamtime at KIT. We acknowledge the provision of beamtime at PETRA III beamline P05 of DESY, a member of the Helmholtz Association (HGF), and support during the beamtime by Hereon team members Fabian Wilde, Julian Moosmann, and Felix Beckmann. Further support was provided in part through the Maxwell computational resources operated at Deutsches Elektronen-Synchrotron DESY, Hamburg, Germany. BEB acknowledges support from the DigiUnit initiative of the Senckenberg Gesellschaft für Naturforschung (2024–2025), and CT is grateful to the support of the Summer Scholarship program from the office of the provost at SUNY Farmingdale (2024). This work is dedicated to the memory of Massimo Olmi, who passed while we were performing the CT data processing.

## AUTHOR CONTRIBUTIONS

B.E.B. Conceptualization; Methodology; Formal Analysis; Investigation; Resources; Data Curation; Writing – Original Draft; Visualization; Project Administration; Funding Acquisition.

A.A.S. Methodology; Software; Formal Analysis; Investigation; Resources; Data Curation; Writing – Original Draft; Visualization; Funding Acquisition.

S.B. Conceptualization; Investigation; Writing – Review & Editing.

J.H. Methodology; Software; Resources; Data Curation; Writing – Review & Editing.

T.v.d.K. Methodology; Software; Resources; Data Curation; Writing – Review & Editing.

D.L. Conceptualization; Methodology; Writing – Review & Editing.

C.T. Conceptualization; Methodology; Investigation; Resources; Writing – Review & Editing; Funding Acquisition.

## STAR METHODS

### Key resources table

| REAGENT or RESOURCE | SOURCE | IDENTIFIER |
| --- | --- | --- |
| Deposited Data |  |  |
| SR-μ-CT data | This paper | **RADAR4KIT XXX**. |
| Voucher specimens (see Table S1 for more detail) | This paper | SMFENT0001921 (BBKIT033), **XXX** (BBKIT256), USNMENT01900055 (BBKIT152), USNMENT01900180 (BBKIT030), USNMENT01900120 (BBKIT044), USNMENT01900217 (BBKIT107), USNMENT01900111 (BBKIT034), **XXX** (BBKIT027), SMFHYM0004682 (BBKIT071), SMFHYM0004681 (BBKIT071), SMFHYM0005629 (BBKIT146), **XXX** (BBKIT071), USNMENT01900426 (BBKIT055), USNMENT01900115 (BBKIT028), USNMENT01900493 (BBKIT256), USNMENT01900494 (BBKIT256), USNMENT01900181 (BBKIT030), USNMENT01818564 (BBKIT093), USNMENT01900295 (BBKIT130), USNMENT01900427 (BBKIT055), **XXX** (DESY smfh_038). |
| Morphometric data | This paper | **Zenodo XXX**. |
| Videographic data | This paper | **Zenodo XXX**. |
| Comparative acceleration data | Booher *et al.* (2021) | https://doi.org/10.5061/dryad.d7wm37q0t |
| Software and Algorithms |  |  |
| MATLAB | MathWorks Inc. |  |
| Adobe Creative Suite | Adobe |  |
| Fiji | ... |  |
| Dragonfly 3D World | Comet Technologies Canada Inc. |  |
| VG Studio | Volume Graphics, Hexagon, Heidelberg Germany |  |
| R (Version 4.2.3) | R Project for Statistical Computing |  |
| R package: ‘car’. | Fox & Weisberg (2019) |  |
| R package: ‘ggplot2’. | Wickham 2016 | ... |
| R package: ‘lmtest’. | Zeileis & Hothorn (2002) | ... |
| R package: ‘sandwich’. | Zeileis *et al.* (2020), Zeileis (2004) | ... |
| R package: ‘dplyr’ (version 1.1.1). | Wickham *et al.* (2023a) | ... |
| R package: ‘ggpubr’ (version 0.6.0). | Kassambara *et al.* (2023) | ... |
| R package: ‘scales’ (version 1.3.0). | Wickham *et al.* (2023b) | ... |
| R package: ‘tidyverse’. | Wickham *et al.* (2019) | ... |
| ... | ... | ... |
| ... | ... | ... |

## RESOURCE AVAILABILITY

### Lead contact

Further information and requests for resources should be directed and will be fulfilled by the lead contact, Brendon E. Boudinot.

### Materials availability

Specimens used for scanning were sourced from the California Department of Food and Agriculture (CDFA), Brendon E. Boudinot research collection (BEBC), and Daniel Tröger research collection (DTC), and are deposited in the Senckenberg Naturmuseum Frankfurt Hymenoptera collection (SMFH). Collection details for the voucher specimens are reported in **Table S1**.

### Data and code availability

The SR-µ-CT, high-speed film, and morphometric data supporting the main findings of this study are available for download at the following link (RADAR4KIT).

**Table S1.**
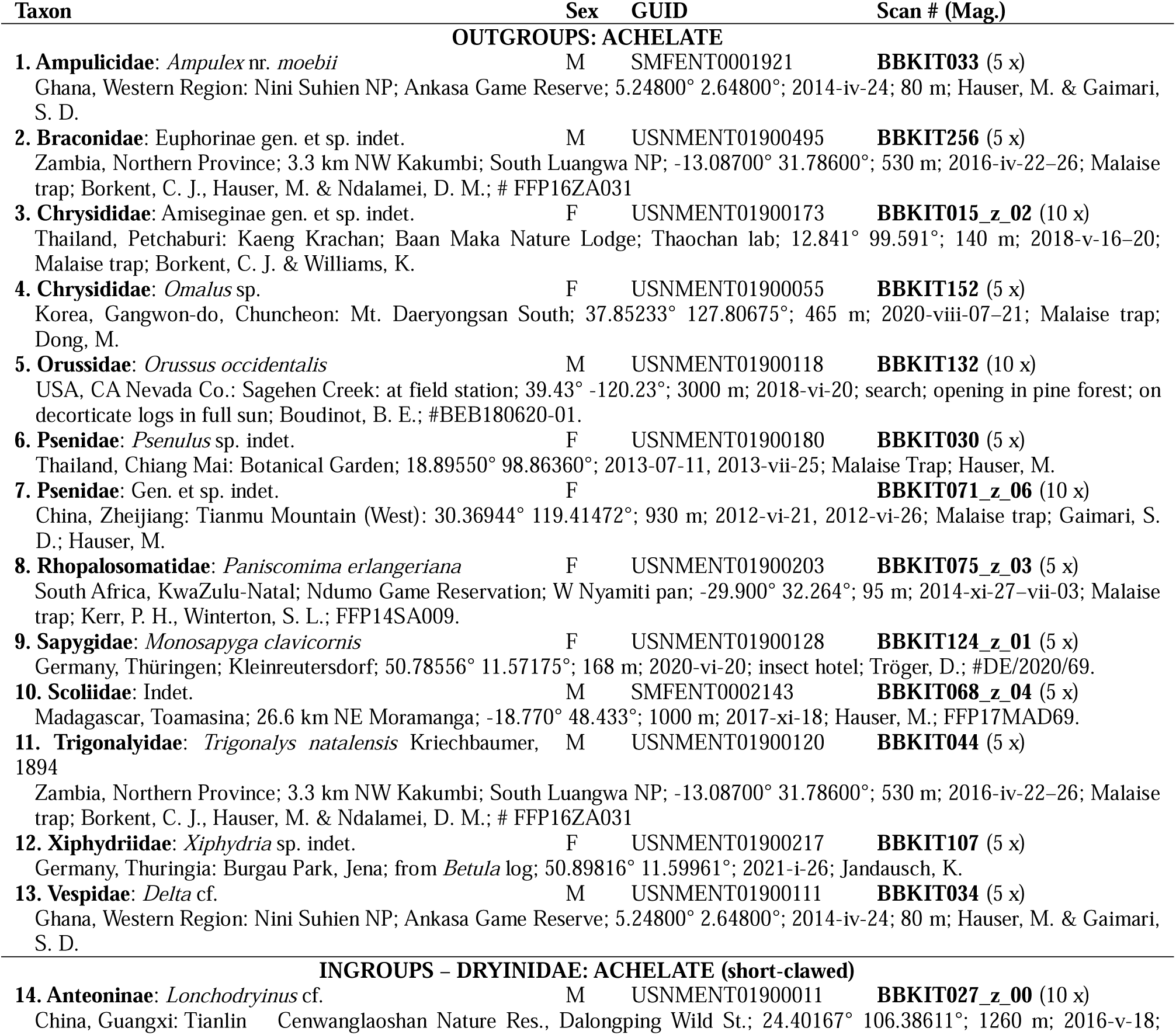
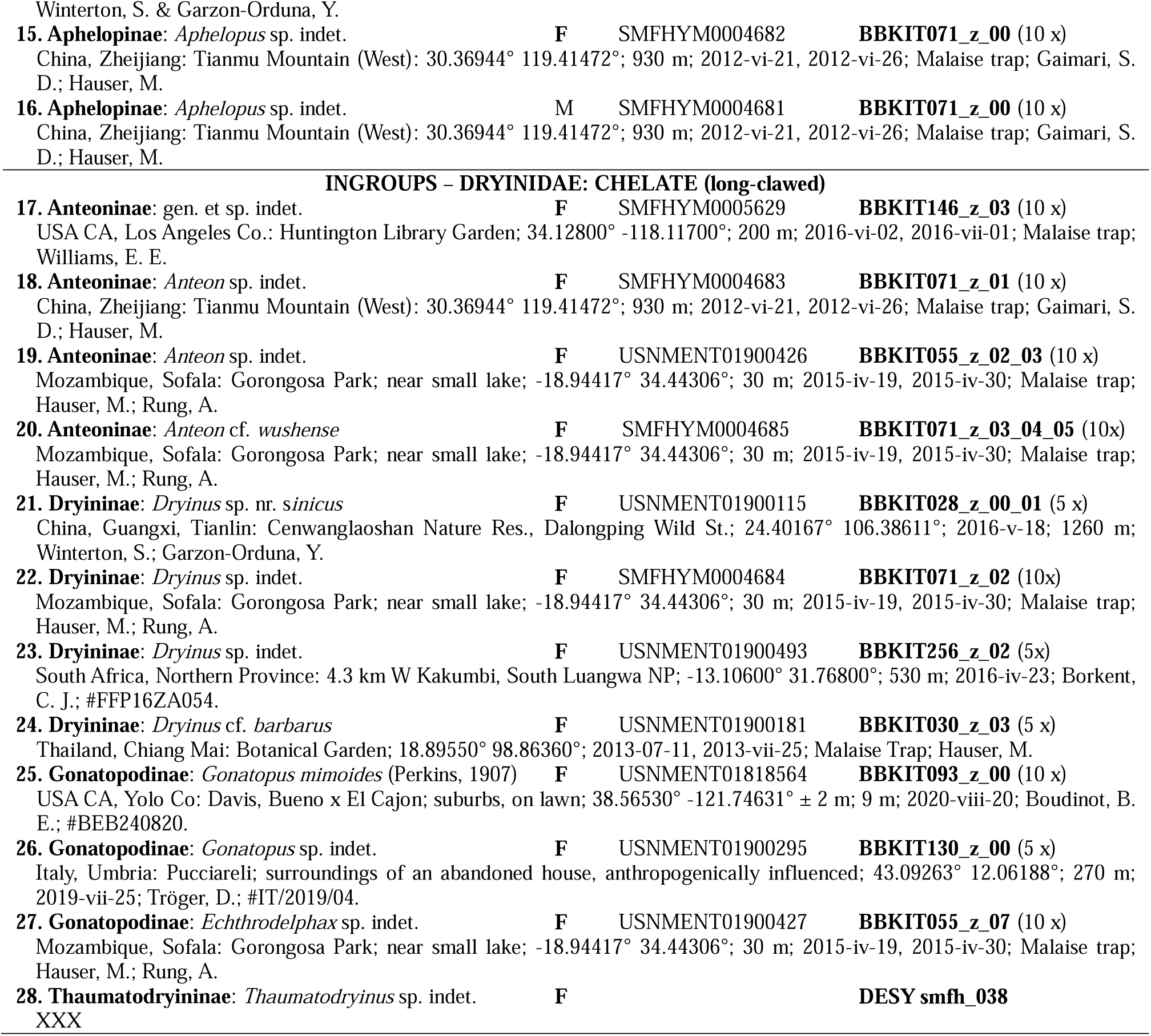
Collection and scan data for the sampled Dryinidae and outgroups.

## EXPERIMENTAL MODEL AND STUDY PARTICIPANT DETAILS

*Not applicable*.

## METHOD DETAILS

### Imaging and reconstruction

#### X-ray scanning

Synchrotron microtomography (SR-µ-CT) for specimens 1–19 (**Table S1**) was performed at the imaging cluster of the KIT Light Source using a parallel polychromatic X-ray beam produced by a 1.5 T bending magnet. The beam was spectrally filtered by 0.5 mm aluminum with a spectrum peak at about 15 keV. We employed a fast indirect detector system, consisting of a 13 µm LSO:Tb scintillator (Cecilia *et al*., 2011), and a diffraction limited optical microscope (Optique Peter, Lentilly, France; Douissard *et al*., 2012) coupled with a 12bit pco.dimax high speed camera with 2016 x 2016 pixels (dos Santos Rolo *et al*., 2014). The specimens were scanned in 95% ethanol. We took 3,000 projections at 70 fps and an optical magnification of 5 and 10 x, resulting in an effective pixel sizes of 2.44 and 1.22 µm. Since the specimen was too large to fit in the vertical field of view, it was scanned in three height steps. The control system concert (Vogelgesang *et al*., 2016) was used for automated data acquisition and online reconstruction of tomographic slices for data quality assurance. The final tomographic 3D reconstructions were performed with tofu (Faragó *et al*., 2022) and included phase retrieval (Paganin *et al*., 2002), ring removal, 8-bit conversion, and blending of phase and absorption 3D reconstructions in order to increase contrast between the background and homogeneous regions, while at the same time highlighting the edges.

Specimen 20 (**Table S1**) was SR-µ-CT scanned at the Imaging Beamline P05 (IBL) (Haibel et al. 2010, Greving et al. 2014, Wilde et al. 2016) operated by the Helmholtz-Zentrum Hereon at the storage ring PETRA III (Deutsches Elektronen Synchrotron, DESY, Hamburg, Germany). A photon energy of 18 keV and a sample to detector distance of 50 mm was used. Projections were recorded using a 20 MP CMOS camera system (Lytaev *et al*. 2014) with an effective pixel size of 0.644 µm. 2501 projections were recorded for each tomographic scan at equal intervals between 0 and π, with an exposure time of 140 ms. The specimen was too large to fit into the field of view in the z-axis, so we scanned overlapping sections and subsequently stitched them together. Tomographic reconstruction was done by applying a transport of intensity phase retrieval and using the filtered back projection algorithm (FBP) implemented in a custom reconstruction pipeline (Moosmann et al. 2014) using MATLAB (Math-Works) and the Astra Toolbox (Palenstijn et al. 2011; van Aarle et al. 2015, 2016). For further processing, the raw projections were binned two times resulting in an effective pixel size of the reconstructed volume of 2.56 µm. For segmentation and visualization, the 32-bit .tif image sequences were converted to 8-bit files and downsampled twofold with Fiji (Schindelin et al. 2012), resulting in an effective pixel (voxel) size of 5.12 µm.

#### 3D Reconstruction

We conducted 3D reconstruction via manual segmentation in the program Dragonfly (Comet Technologies Canada Inc., Montréal, Québec). We used a multi-slice brush to manually label regions of interest (ROIs) in a multi-ROI with a variably set threshold. To export TIFF stacks, we followed the steps of Boudinot *et al*. (2024), including ROI extraction and inversion, dataset duplication and masking, and finally export. For volumetric rendering, we imported these image stacks into VG Studio (Volume Graphics GmbH, Heidelberg, Germany) and rendered using Phong shading and two light sources; clipping planes were used on individual objects for rendering digital dissection. 3D prints were made to physically assess patterns of joint interaction also following the protocol of Boudinot *et al*. (2024).

#### Figure composition

Figures were composed using Adobe Illustrator. Where light / dark or other tone adjustments were necessary, and would not affect inferences, Adobe Photoshop was used.

### High-speed filming

High-speed video sequences of strikes were captured with a Phantom VEO 1310 camera through a Venus Optics Laowa 60mm f/2.8 2x Ultra-Macro lens with the aperture set at f/11. The sequences were recorded at a rate of 6,000 frames per second with an 83.33 µs exposure. The scene was illuminated from behind with constant diffused light and above from a by a high-intensity 12,000-lumen LED array at 50% power (Visual Instrumentation Corporation). A live *Gonatopus* sp. wasp and early instar planthopper (cf. Acaloniidae) were placed on a filming platform which was a flat piece of acrylic (6 cm x 1.9 cm) covered in a general-purpose beige masking tape (Duck® brand) to provide a uniformly textured material while maintaining a flat substrate. The platform was manually moved from below so that the insects remained in the focal plane while they freely explored the platform. The camera was set to record using a post-trigger which allowed for capturing a maximum of 1.65 seconds of action before the trigger point. This was enough time to capture the entire behavioral sequence of a strike, from first approach to retracting a caught prey item to the body.

### Morphological terminology

The general morphological terminology for the leg follows Snodgrass (1935), with revisions for tarsal anatomical terms and concepts from Beutel *et al*. (2020) and musculature from Aibekova *et al*. (2022) and its recent update (Aibekova *et al*. 2025). The terms “chela”, “chelate”, and “achelate” are conventionally used in the Dryinidae literature (*e.g.*, Olmi 1984) to refer to the anterior pretarsal claw (as newly determined here) or to the condition of having an elongated anterior pretarsal claw or not. Insect limbs have historically been treated as conservative, groundplan-constrained systems (e.g., XX_Snodgrass-1935), with little attention paid to their comparative internal variation. The leg system definition provided in the results section, therefore, should be used as a general reference or comparison point for insect legs.

## QUANTIFICATION AND STATISTICAL ANALYSIS

### Overview

We quantified (1) the functional geometry of the claw actuation system from our SR-µ-CT data and (2) the rate of claw closure from our high-speed videography. The CT-based quantifications and analysis are presented first, followed by the work calculations.

### Geometrical (Anatomical) Measurements

Muscle measurements were taken in Dragonfly using volumetric segmentation or the ruler and path tools in cross-sectional view. For each linear measurement, the virtual reregistration tool was used to establish exact cross, frontal, and sagittal views.

#### Muscles

##### Volume

To quantify muscle volume, a threshold was set that (a) allowed for the capture of all pixels attributable to musculature at the lower end, and (b) the exclusion of sclerite and apodeme at the upper end. With such a threshold, the multi-slice paintbrush tool was used to manually assign pixels to the Ifpm1 and Itpm1 muscles. Where the muscles attached to the cuticle, the surface between the muscle and cuticle was also included. To ensure that cuticle was outside of the selectable threshold range, the lookup table (LUT) was set to “greyscale” with black being empty and white representing x-ray dense material. The muscle Itpm1 had very few fibers and was excluded from downstream analysis. The specimen of *Dryinus* cf. *barbarus* (“*Mesodryinus*”) was excluded as the preservation was inadequate, having apparently dried out at some point during its archival history. All dryinid specimens that were used for volumetric analysis had their chelae closed, hence were comparable with respect to the length of muscle fascicles and their pennation angles.

##### Fiber length

Individual fibers were identified in the 2D viewer of Dragonfly by tracing their down length using the virtual re-registration crosshairs, then measured using the line ruler tool. At least 10 measurements were made for the *M. profemuro-pretarsalis* (L(Ifpm1), Fig. S1) and as many as possible were made for *M. protibialo-pretarsalis* (Itpm1). These measurements were averaged.

##### Fiber pennation angle

For each muscle bundle, fiber pennation angle (θ, °(Ifpm1), Fig. S1) was averaged across ≥ 10 measurements using the angle tool, unless the muscle bundle had < 10 fibers, in which case as many angles were taken as possible. The longitudinal axis of the pretarsal apodeme was approximated as the line of action, and the vertex of the angle set at the detectable tendonal attachment of the fiber.

**Fig. S1.**
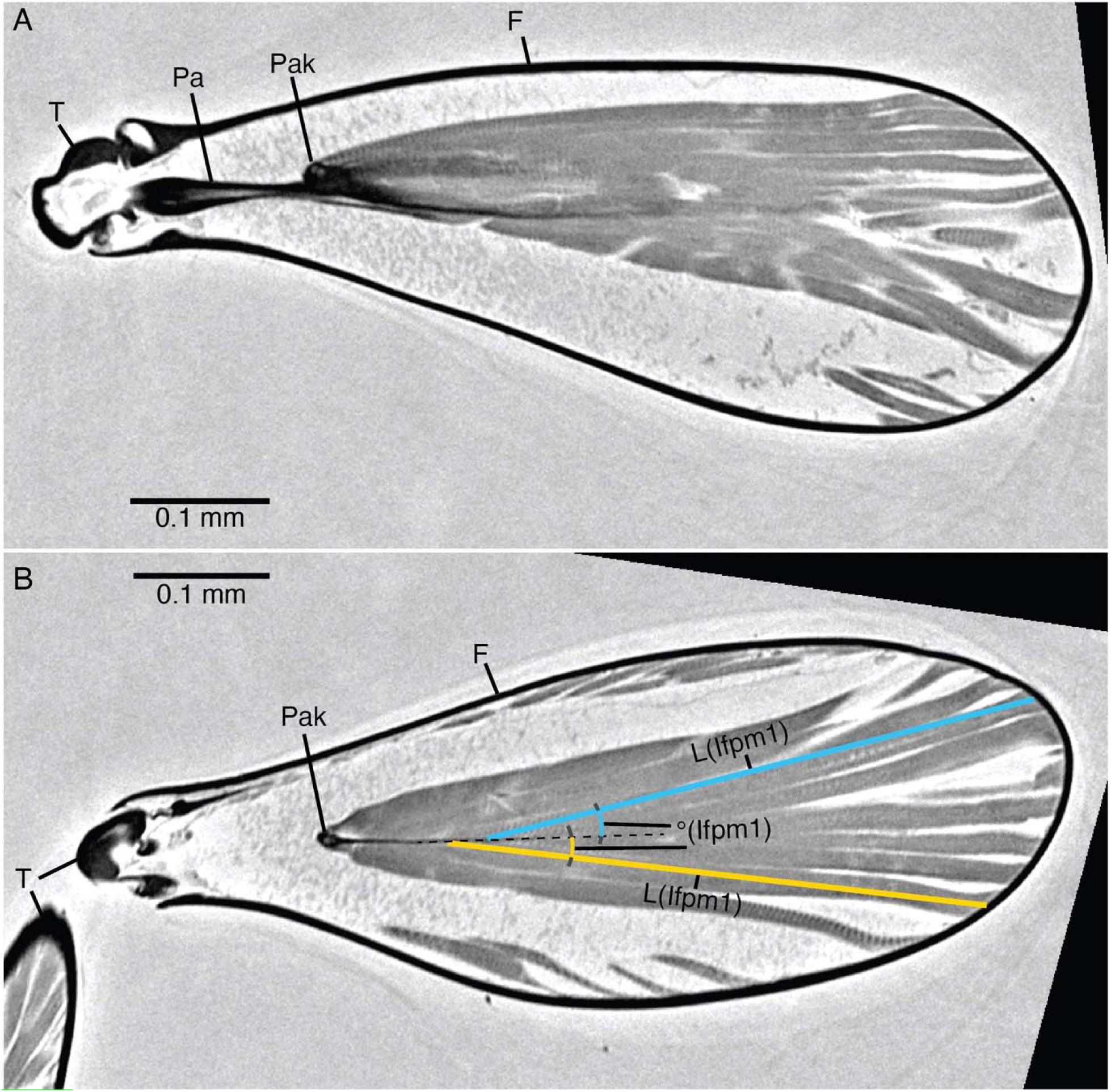
Demonstration of muscle fiber length and angle measurement (Anteoninae, bb146_z_03). **A**: Sagittal view, through keel of pretarsal apodeme. **B**: Frontal view, through keel of pretarsal apodeme. **Abbreviations**: F = femur; L(Ifpm1) = length of Ifpm1 muscle fiber; Pa = pretarsal apodeme; Pak = pretarsal apodeme keel; T = tibia; °(Ifpm1) = pennation angle of Ifpm1 muscle fiber.

#### Characteristic lengths

##### Pretarsal apodeme length

Because the pretarsal apodeme bends along its length both due to preservation and curvature in the leg joints, it was necessary to estimate its length with a combination of multiple tools. Inside the femur and tibia, the ruler tool was used to approximate the length in linear steps; measurements within each of these segments were taken at least twice and averaged. In the joints, the path tool was used to trace along the curve of the apodeme, with multiple measurements per bend averaged.

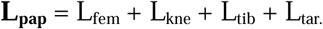

##### Unguitractor plate length

In cross-section, the unguitractor plate is identifiable as a thickening that has discrete proximal and distal boundaries, as well as an internal longitudinal hollow. The length of the plate was measured along the length of the hollow from the one end to the other (Fig. S2D–F).

**Fig. S2.**
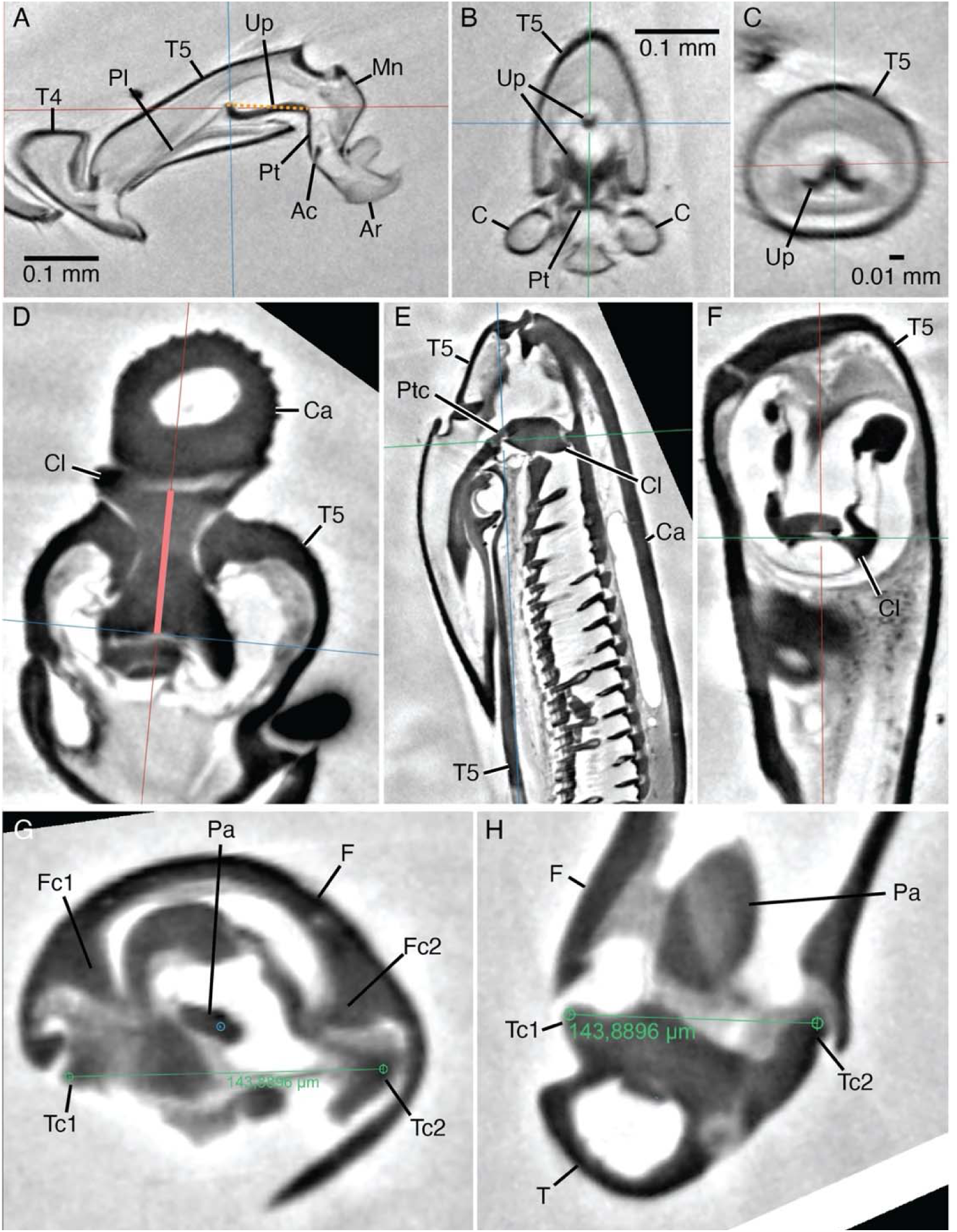
(**A–C**) Length measurement for the unguitractor plate in achelate taxa (specimen: *Trigonalys* sp.). (**A**) Sagittal section. (**B**) Frontal section. (**C**) Cross sectional. (**D–F**) Length measurement for the Olminator ligament (specimen: *Drynius* cf. *sinicus*); each panel is of the same moment of viewing but not to scale; the thick red line in D was used to measure L_os_. (**A**) Frontal section. (**B**) Sagittal section. (**C**) Cross section. (**G, H**) The “knee width” measurement (L_k_), which is taken from across the width of the protibia (T) between the proximal protibial condyles (Tc1, Tc2), as exemplified by *Dryinus* cf. *barbarus* (bb030). (**G**) Cross sectional view. (**H**) Frontal view. **Abbreviations**: Ac = arculus; Ar = arolium; C = pretarsal claw; Ca = anterior pretarsal claw (= chela); Cl = pretarsal claw ligament; F = femur; Fc1, Fc2 = distal femoral condyles; Mn = manubrium; Os = Olminator ligament; Pa = pretarsal apodeme; Pl = pretarsal apodeme; Pt = planta; Ptc = connecting ligament of pretarsal apodeme; T = tibia; T4 = fourth tarsomere; T5 = fifth tarsomere; Tc1, Tc2 = proximal tibial condyles. Up = unguitractor plate.

##### Proximal unguitractor apodeme length

Proximad the unguitractor plate, the unguitractor apodeme opens to the outside ventrad the pretarsal complex. The length of this portion of the apodeme was measured along the length of its hollow to its proximal aperture using both the line and path tools in slice view, which necessitated carefully following the apodeme along its length and resetting the viewing planes using the virtual reregistration axes of Dragonfly.

##### Femorotibial knee width

In order to control for body size, a characteristic length is needed. Because the scaling rules of the three tagmata of hymenopteran bodies have not been established, we chose to use the width of the femorotibial “knee” width, or proximal protibial width, as a candidate for characteristic length. Knee width (**W_tib_**) was measured as the maximum external diameter of the proximal joint of the protibia ventrad the femoral condyles. We expect that variation of this width among Dryinidae will be constrained by natural selection, whereas the shape of the head in this family, for example, is known to vary considerably and may not provide a robust proxy for body size (Olmi 1984). As expected for a proxy of body size, we found that **W_tib_** scales isometrically with pretarsal apodeme length (**L_pap_**) (standardized major axis [SMA] [XX-Warton-et-al.-2012], *log-transformed, single-group test: H0, slope not different from 1: r=0.030, df=17, p=0.903*). We therefore used **W_tib_** as our characteristic length for the claw system, as it was easier to measure accurately than **L_pap_**.

#### Claw clamp mechanism

##### Mechanical advantage

The mechanical advantage of the anterior pretarsal claw was calculated by dividing the length of the inlever (***a***) by the length of the outlever (***b***).

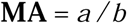

The in- and outlever measurements were taken in a single plane set to the sagittal section of the claw (Fig. S3). Because this plane is perpendicular to the articulation between the claw and fifth tarsomere, the articulation approximates a normal vector, hence is the axis around which the claw rotates. This articulation spans the diameter of the claw, which is proportionally wider than other Hymenoptera. The central point of the anterior claw fulcrum was approximated as the midpoint of a line set along the length of the hinge between the fifth tarsomere and the claw. The endpoint of the inlever was set as the middle apex of the proximoventral rim of the anterior claw (*a* in Fig. S3B), where the olminator sclerite would reasonably make contact during motion. The endpoint of the outlever was set at the apex of the anterior claw (*b* in Fig. S3B). For the outgroup taxon *Trigonalys*, which was bidentate without an obvious single terminus, the outlever measurements to both apices were averaged for.

**Fig. S3.**
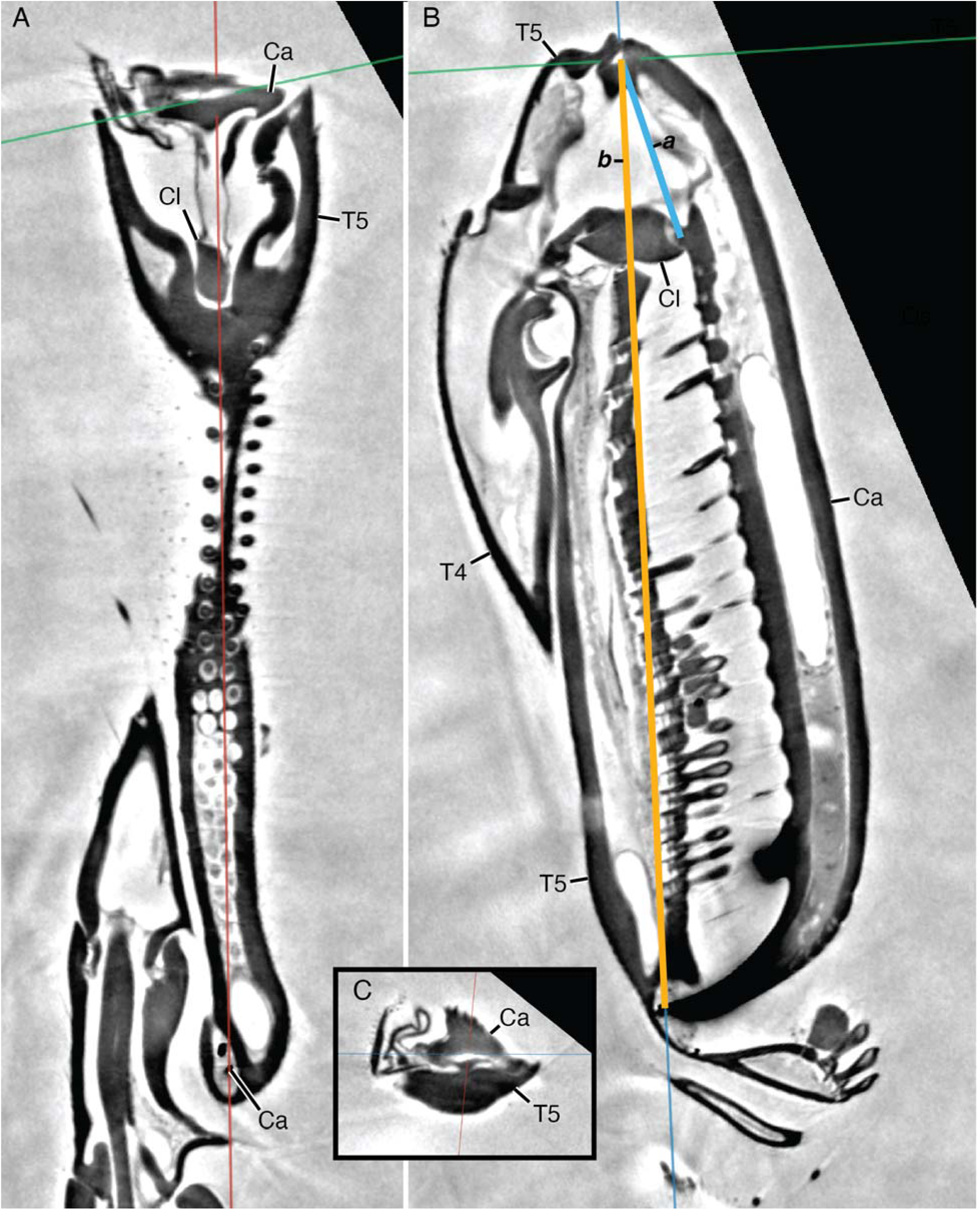
Measurements required to calculate MA (mechanical advantage) (specimen: *Dryinus* cf. *sinicus*). Each panel is of the same moment of viewing but not to scale. (**A**) Frontal section. (**B**) Sagittal section. (**C**) Cross section. **Abbreviations**: *a* = inlever length; *b* = outlever length; Ca = anterior pretarsal claw (= chela); Cl = pretarsal claw ligament; Os = Olminator ligament; T4 = fourth tarsomere; T5 = fifth tarsomere.

##### Expected force output capacity

To estimate the geared force output capacity of the claw actuation system (**A_eff_**), we multiplied the physiological cross-sectional area (PCSA) of the femoral-tibial pretarsal claw closer muscle (Ifpm1) by the mechanical advantage (MA) of the anterior claw (Püffel *et al*. 2021):

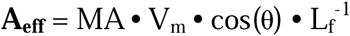

It was not possible to measure the physiological strain of the muscle as all SR-µ-CT-scanned specimens were preserved.

**Fig. S4.**
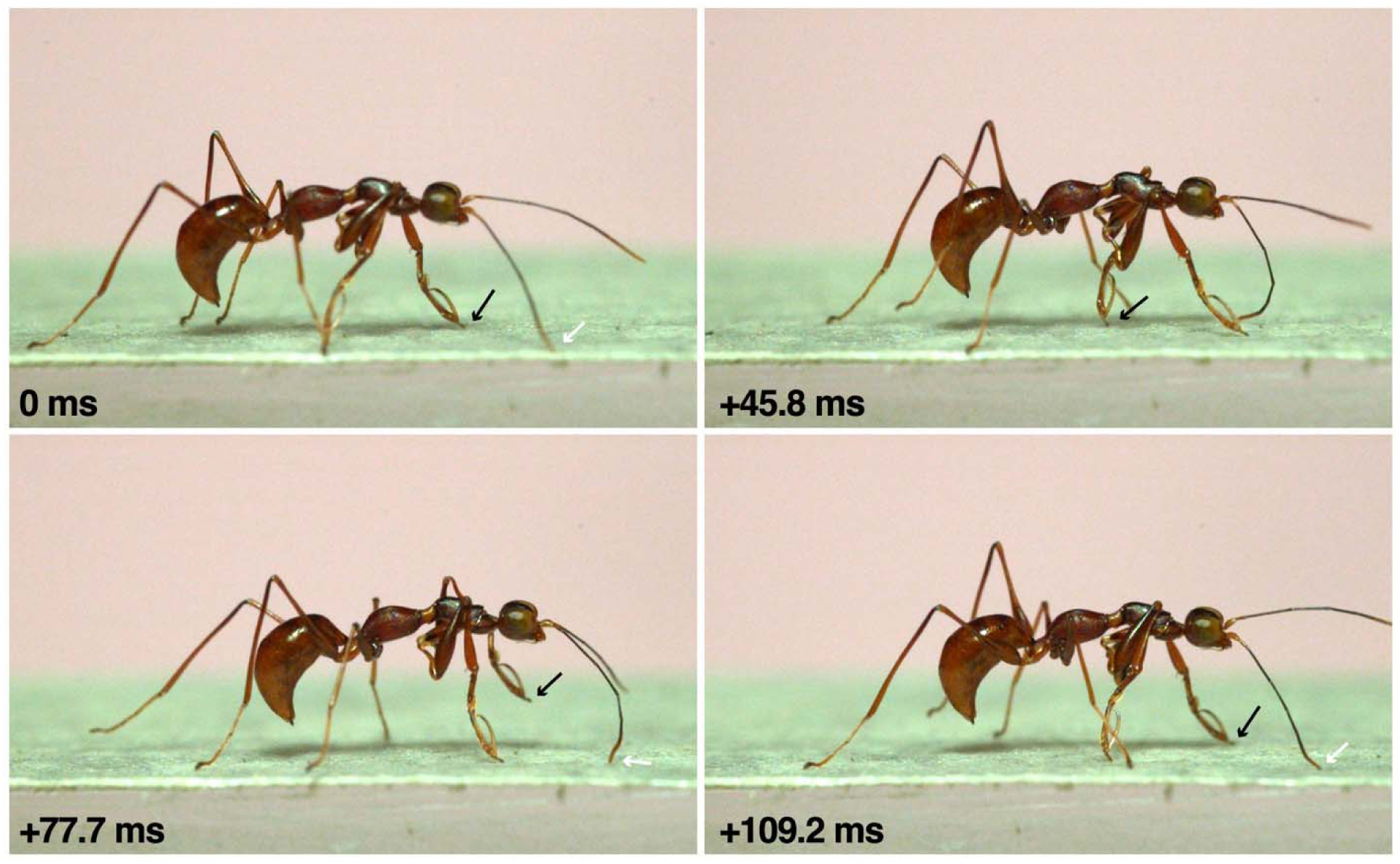
Walking cycle of *Gonatopus* sp., showing derived aroliograde locomotion of prothoracic leg, which compensates for the extreme modification of the anterior pretarsal claw.

### Statistical Tests and Physical Calculations

#### Data exclusion

Due to the poor preservation of the tibial musculature across the sampled Hymenoptera, the muscle geometries could not be measured for all specimens. We have observed that this is a general preservational issue based on a large quantity of scans. This issue is most likely due to gas exchange across the tibial trachea, which can often be observed to fill a large volume of the tibia in specimens freshly killed in ethanol. In other specimens, the femoral musculature was poorly preserved. These cases were most likely due to the variable kill methods and preservation histories, as the specimens were collected from the field. No specimens were excluded from analyses involving cuticular morphometrics.

#### Statistical analysis of allometric relationships (OLS)

We quantified allometric relationships among claw biomechanical variables in Dryinidae using ordinary least-squares (OLS) regression in R (R Core Team 2023). To determine primary linear size measure (characteristic length), we regressed pretarsal apodeme length (**L_pap_**, ‘L_apod’, in mm) on tibial width (**W_tib_**, ‘W_tib’, in mm) using both untransformed and log_10_-transformed variables and inspected residuals and Breusch–Pagan tests for heteroscedasticity. Based on the high correlation between the two variables and the ease of measuring tibial width, we chose **W_tib_** as the characteristic length.

For the main analyses, we log_10_-transformed all morphometric and biomechanical variables prior to modeling: inlever length (***a***, ‘a_inlever’), outlever length (***b***, ‘b_outlever’), mechanical advantage (**MA**, ‘MA’), and expected force output (**A_eff_**, ‘A_eff’, or PCSA • MA). One specimen of chelate Dryinidae, *Anteon* cf. *wushense*, was appears to be an outlier with a proportionally long inlever; it had no statistical effect.

All log-log allometries were fit with OLS using models of the general form

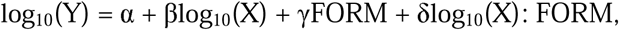

where Y was one of {‘a_inlever’, ‘b_outlever’, ‘MA’, ‘A_eff’}, X was either ‘W_tib’ or ‘a_inlever’, and FORM is a fixed factor with two levels (‘achelate’, ‘chelate’). For each relationship we first fit a full interaction model (size • FORM) and then a reduced additive model without the interaction term (common slope, different intercepts). Model summaries were obtained with ‘lm’, and nested models were compared with F-tests via ANOVA.

To assess homoscedasticity, we examined residual-versus-fitted plots and performed Breusch-Pagan tests using ‘bptest’ (‘lmtest’). Where the Breusch-Pagan test indicated heteroscedasticity (*e.g.*, several models for ‘b_outlever’ versus ‘a_inlever’, ‘MA’ versus ‘W_tib’, and ‘A_eff’ versus ‘W_tib’), we recalculated heteroscedasticity-consistent standard errors using the sandwich estimator (‘vcovHC’ from the R package ‘sandwich’) and ‘coeftest’ (‘lmtest’) for four variants: HC1, HC2, HC3, and HC4. We treated HC3 and HC4 as the most conservative; significance and confidence in slope and intercept estimates were checked across these estimators.

We tested isometry and other hypotheses about slopes using linear constraints on the regression coefficients. Specifically, we used ‘linearHypothesis’ (‘car’) to test:

**Isometry**: **H_0_**: β = 1 for the size exponent (*e.g.*, ‘log_W_tib’ = 1 for size-trait scaling, or ‘log_a_inlever’ = 1 for outlever versus inlever),
**No scaling**: **H_0_**: β = 0 (*e.g*., for ‘MA’ vs ‘W_tib’),
**Group-specific slopes**: linear combinations such as **H_0_**: β_Wtib + β_Wtib:FORMChelate = 1 or 0 for the chelate slope, and analogous tests for achelate slopes.

Where one-sided alternatives were biologically motivated (*e.g.*, testing whether chelate slopes exceeded isometry), we computed manual one-sided t-tests by combining coefficients and their covariance from ‘vcov’ model and evaluating the resulting t-statistic with the appropriate residual degrees of freedom.

Visualization of fitted allometries was performed with ‘ggplot2’. For regression displays, we plotted log_10_-transformed data with OLS lines (‘geom_smooth(method = "lm")’) and 95% confidence bands. For the figures, axes were labeled in raw mm while the underlying coordinates remained log_10_-transformed: this was done by specifying log-transformed breaks and using their inverse (raw mm) as axis labels (scale_x_continuous / scale_y_continuous with breaks = log_10_(mm_breaks), labels = mm_breaks). To compare inlever and outlever scaling directly, we reshaped the data to long format (pivot_longer) and plotted both levers against log_10_(W_tib) in a single panel, using different symbols for lever type and different colors for FORM, with separate OLS lines and confidence intervals for each group/lever combination.

### Power density estimation

We observed that claw closure for both the left and right claws our clearest high speed videographic sequence, represented in Fig. 1I, J lasted no more than two frames (0.00016667 s). To conservatively estimate the power density with this information, we used measurements from a similarly sized congeneric species (*Gonatopus*, sample bb130). We measured the volume of claw cuticle from our segmented data (0.00026523 mm^3^), hence a reasonable mass of 2.6523E-10 kg. The center of mass of the claw cuticle was calculated using the “center of mass” tool in Dragonfly. We then measured the length of the claw from its axis of rotation to the CoM (0.0001694 m). Assuming a conservative arc of closure at 90° and using the CoM coordinate measurement, we estimate that the angular velocity was at least 9424.6 radians / s, with the CoM traveling an arc length of 0.000266093 m, resulting in an average linear velocity of 1.59653 m /s, with 3.8022E-10 J of kinetic energy and 2.02809E-06 W of power. Given that the volume of the closer muscle was measured to be 2.36827E-12 m^3^, hence with an estimated mass of 2.51037E-09 kg, the power density would be ∼807.9 W / kg.

